# Short communication: Validation of a novel milk progesterone-based tool to monitor luteolysis in dairy cows using cost-effective, on-farm measured data

**DOI:** 10.1101/526061

**Authors:** Ines Adriaens, Wouter Saeys, Katleen Geerinckx, Bart De Ketelaere, Ben Aernouts

## Abstract

**INTERPRETATIVE SUMMARY. Validation of luteolysis monitoring tool. Adriaens.:** Monitoring of milk progesterone allows to detect luteolysis preceding a dairy cow’s estrus. In this research, two algorithms were compared using on-farm measured milk progesterone data with a variable, cost-effective sampling rate. The first was the state of the art for progesterone monitoring, a smoothing multiprocess Kalman filter compared with a fixed threshold. The second algorithm was the recently developed progesterone-based monitoring algorithm using synergistic control. Both algorithms detected approximately the same amount of luteolyses, but the second gave the alerts on average 20 hours earlier. This proves that this algorithm can operate reliably in an on-farm setting.

---

The Progesterone (**P4**) monitoring algorithm using Synergistic control (**PMASC**) uses the luteal dynamics to identify fertility events in dairy cows (Adriaens et al., 2017, 2018a). Hereto, PMASC employs a combination of mathematical functions describing the increasing and decreasing P4 concentrations during the development and regression of the corpus luteum and a statistical control chart which allows to identify luteolysis. The mathematical model combines sigmoidal functions from which the cycle characteristics can be calculated. Both the moment at which luteolysis is detected and confirmed by PMASC, as well as the model features themselves can be used to inform the farmer on the fertility status of the cows.

Until now, the PMASC methodology was designed and optimized using milk P4 data obtained through experimental trials, collected at maximum frequency (e.g. every milking) and analyzed *post-hoc* using the state-of-the-art laboratory ELISA technique. Moreover, this method was not yet challenged on data sampled at reduced frequency to show its possible cost-effectiveness, nor was it shown that PMASC can work on data measured with an on-line device proving its value as an on-farm monitoring algorithm. The objective of this study was to show the capability of PMASC to work in an on-farm setting, identifying luteolysis using smart-sampled data and analyzed via an on-farm lateral flow immunoassay (**LFIA**). Moreover, we compared the PMASC algorithm with the state of the art for P4 monitoring on farm, i.e. smoothing the raw P4 values, combined with a fixed threshold for detection of luteolysis and alerts.

Datasets of on-farm measured milk P4 were collected on two farms in Flanders, Belgium. The first dataset (dataset 1) was obtained from an experimental research farm between March 2017 and January 2019. On this farm, the herd was split up in two groups, each consisting of on average 55 lactating dairy cows. Both groups were milked with an automated milking system (VMS, Delaval, Tumba, Sweden). The second dataset (dataset 2) was collected on a commercial dairy farm between November 2016 and December 2018. The herd consisted of on average 250 animals, milked by four automated milking systems (VMS, Delaval, Tumba, Sweden). On both farms, animals were fed a mixed ration of grass and corn silage, supplemented with concentrates through automated concentrate feeders and in the milking robot.

For the P4 analysis, milk samples were taken automatically by the sampling unit integrated in the Herd Navigator^TM^ (Delaval, Tumba, Sweden; Lattec, Hillerød, Denmark), and the milk P4 concentration was determined by the built-in LFIA milk analyzer. More details on the working principle of this device can be found in Blom and Ridder, (2010) and Friggens et al., (2007). The samples for P4 analysis were taken with a variable sampling frequency, of which the concept has been described by Friggens and Chagunda, (2005). Basically, the time between samples is determined by the days in milk, the likelihood of luteolysis, the P4 concentration of the previous sample, the performed inseminations and their timing, the likelihood of pregnancy, and the probability of prolonged postpartum anestrus or cysts. As the consumables are an important cost for P4 monitoring on farm, this sampling scheme ensures that the necessary fertility information can be gathered on crucial moments with a minimum number of samples.

Dataset 1 consisted of 10,791 P4 measurements originating from 144 lactations for 104 unique cows (parity 1 to 6, 2.15±1.16, average milk production per day 36.47 ± 12.66 kg, mean ± standard deviation). The second dataset consisted of 36,645 P4 measurements, originating from 460 lactations and 342 unique cows, with parities between 1 and 8 (2.3±1.26, average milk production per day 34.24±10.10 kg).

The state of the art (milk) P4 monitoring in dairy cows builds on a multiprocess Kalman filter (**MPKF**) to smooth the profiles and reduce the effect of potential outliers (Korsgaard and Løvendahl, 2002; Friggens and Chagunda, 2005; Friggens et al., 2007). When the smoothed P4 value undercuts a 5 ng/mL threshold (**T**), after that it exceeded 5 ng/mL twice, luteolysis was deemed confirmed. This algorithm, further referred to as MPKF+T, was implemented and used to identify the reference for luteolysis which was then used to benchmark the performance of our PMASC algorithm in terms of detecting the luteolysis events.

Progesterone profiles (i.e. the P4 time series of one lactation) with less than 20 measurements in total (e.g. because they just started at the moment of data collection), or less than 5 measurements below or above 5 ng/mL were discarded. Also, P4 data following a gap in the P4 data of at least 20 days were excluded. Accordingly, 719 and 5322 measurements originating from 47 and 317 lactations were deleted from each dataset respectively, leaving 10,072 and 31,213 measurements from 133 and 409 lactations belonging to 98 and 316 unique cows. The characteristics of the remaining profiles and related P4 data are summarized in Table 1.

**Table 1.** Summary of the datasets of farm 1 and farm 2

| Dataset | No. of cows | No. of profiles | MY<sup>1</sup> (kg) | MI<sup>2</sup> (h) | P4<sup>3</sup> |  |
| --- | --- | --- | --- | --- | --- | --- |
|  |  |  |  |  | Concentration (ng/mL) | No. of meas. per profile |
| Dataset 1 | 98 | 133 | 9.98 – 65.5<br>(36.47 ± 12.66) | 5.6 - 29.7<br>(7.9 ± 2.62) | 0.03-27.98<br>(14.71±10.23) | 23 - 168 (75.7 ± 29.97) |
| Dataset 2 | 321 | 421 | 10.1-66.4<br>(32.0±11.7) | 5.0-29.05<br>(9.36±3.19) | 0.01-27.99<br>(15.36±9.69) | 21-232<br>(80.2±35.39) |

<sup>1</sup> Daily milk yield in kg; <sup>2</sup> Average milking interval in hours; <sup>3</sup>P4 milk progesterone;

Adriaens et al., (2018a) describes the original PMASC to process time series of P4 measurements taken each milking, two to four times a day. To avoid false alerts caused by coincidentally low P4 values (e.g. outliers), a first evidence of luteolysis (low P4) identified by the statistical control chart needed to be confirmed by two additional successive measurements before a luteolysis alert was raised by PMASC. However, in the datasets employed in this study, the P4 concentration was not available for every milking as the number of and moment of samplings depended on the algorithm implemented in the analyzer. For example, when an upcoming estrus was confirmed by this algorithm, no new milk samples were analyzed in the following 3 to 6 days. In case of a steep drop in milk P4 in combination with very low absolute P4 concentrations (< 5 ng/mL), this often resulted in less than the three consecutive out-of-control measurements available to confirm luteolysis with PMASC. Therefore, the main decision rule of PMASC needed to be adjusted in function of these new datasets. In previous studies on the timing between luteolysis and ovulation time (Roelofs et al., 2006; Adriaens et al., 2018b), it was found that ovulation followed about 60 to 80 hours after the onset of luteolysis. To have the largest chance on successful conception, insemination should take place 24 to 12 hours before ovulation (Roelofs, 2005). Based on this information, we evaluated all the samples collected within 36 hours following an out-of-control measurement to confirm or object luteolysis. More concretely, following rule was implemented: if more than half of the measurements within this 36h window were also out of control, luteolysis was assumed confirmed, an alert was raised and a new cycle started. In the other case, the out-of-control measurement was considered an outlier. In practice, this would ensure that luteolysis has taken place, while there would have been still sufficient time left to arrange the insemination. Moreover, this rule gives some flexibility to deal with missing samples and outliers. In future studies when the sampling frequency can be defined based on PMASC, other rules can be implemented.

Next, PMASC was applied to the time series of P4 values of each lactation to evaluate its performance when in this cost-effective setting controlled by the device’s algorithm. The number of and the timing of the alerts generated by PMASC were compared to those obtained using the MPKF+T algorithm. Also, the number of measurements per detected cycle was calculated and the cycle characteristics were summarized.

An overview of the results is given in Table 2. Dataset 1 consisted of 133 P4 profiles, with an average sampling rate of 0.48±0.07 measurements per day in the period between the first and the last measurement. This corresponds to one measurement each 2.07 days. For dataset 2, the 409 profiles had an average sampling rate of 0.50±0.07 measurements per day, corresponding to one sample per 1.98 days. The MPKF+T algorithm gave 427 and 1416 luteolysis alerts (i.e. smoothed time series undercutting the fixed threshold of 5 ng/mL), respectively for dataset 1 and 2, while PMASC gave 420 and 1411 alerts. Alerts within 5 days of each other were taken as the same luteolysis. Based on this timing, 25 (5.8%) and 54 (3.81%) of the alerts of MPKF+T could not be matched with one of PMASC, and 18 (4.3%) and 49 (3.47%) of the alerts of PMASC could not be matched with one given by MPKF+T. These unmatched alerts could be attributed to (1) a sampling frequency too low to confirm the out-of-control samples, for example due to quite low luteal phase P4 values, after which only one sample sufficed to generate an alert using MPKF+T. This was the case for respectively 58% and 42% of the unmatched alerts for dataset 1 and 2; (2) high follicular phase P4 values, resulting in enough evidence for PMASC to generate an alert, while MPKF+T did not yet undercut the threshold value. In this case, sometimes the MPKF+T alert followed later (but outside the time window for matching i.e. 5 days. This was the case for 42% and 58% of the unmatched alerts in dataset 1 and 2, respectively. In the first case, the smart sampling algorithm does not take a new confirmatory sample, while we set PMASC to evaluate at least two samples per estrus, and PMASC thus failed to give an alert. When the sampling is based on decision rules implemented in PMASC, outside the scope of this research, also these luteolyses can be detected by PMASC. The intermediate values causing additional PMASC alerts in the second case (2), are mainly due to the calibration technique of the online measurement device, as also discussed in Adriaens et al., (2019).

**Table 2.**
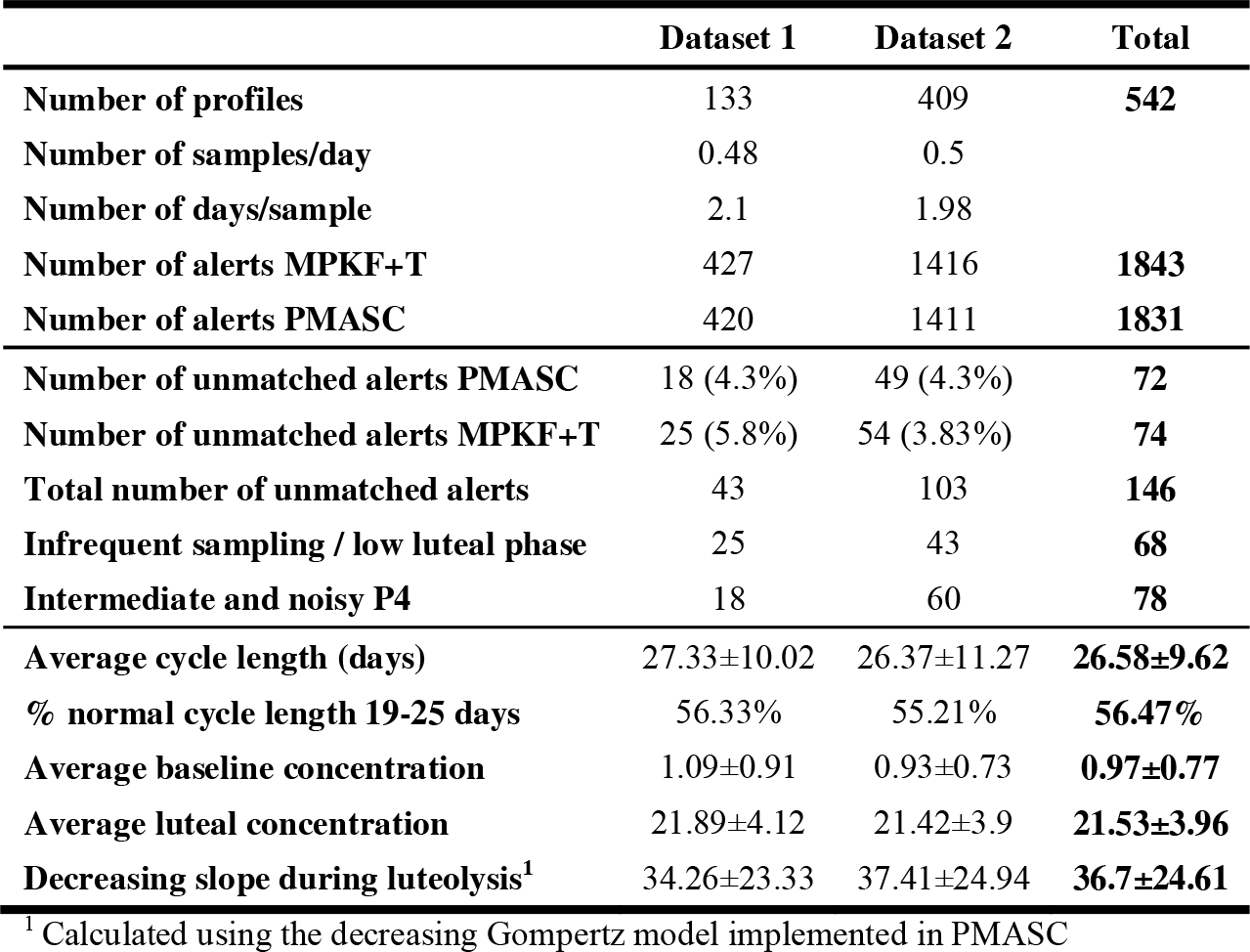
Overview of the detected luteolysis by each algorithm and the corresponding cycle characteristics

An example of a P4 profile and the corresponding alerts is shown in Figure 1. For the first cycle, no alert was generated with PMASC. However, the sample for which the smoothed value is below the 5 ng/mL threshold was also out-of-control in PMASC, but the next sample was only taken 6 days later. As such, luteolysis was not detected by PMASC because of the lack of confirmatory samples out-of-control within 36 hours after the first out-of-control measurement. The final PMASC models for which the detected luteolysis matches with the MPKF+T algorithm, while the MPKF smoothed data are given in dark blue. The threshold of 5 ng/mL P4 is plotted in green. As seen in the figure, the first sample detected as out of control by PMASC is often not yet below the 5 ng/mL threshold, and the model characteristics can be used to estimate luteolysis reliably and independent of both the milking frequency and the absolute P4 values. The time lag between the drop in P4 during luteolysis and the smoothed MPKF value undercutting the threshold is shown to be dependent of sampling rate and the absolute P4 values (yellow box).

**Figure 1.**
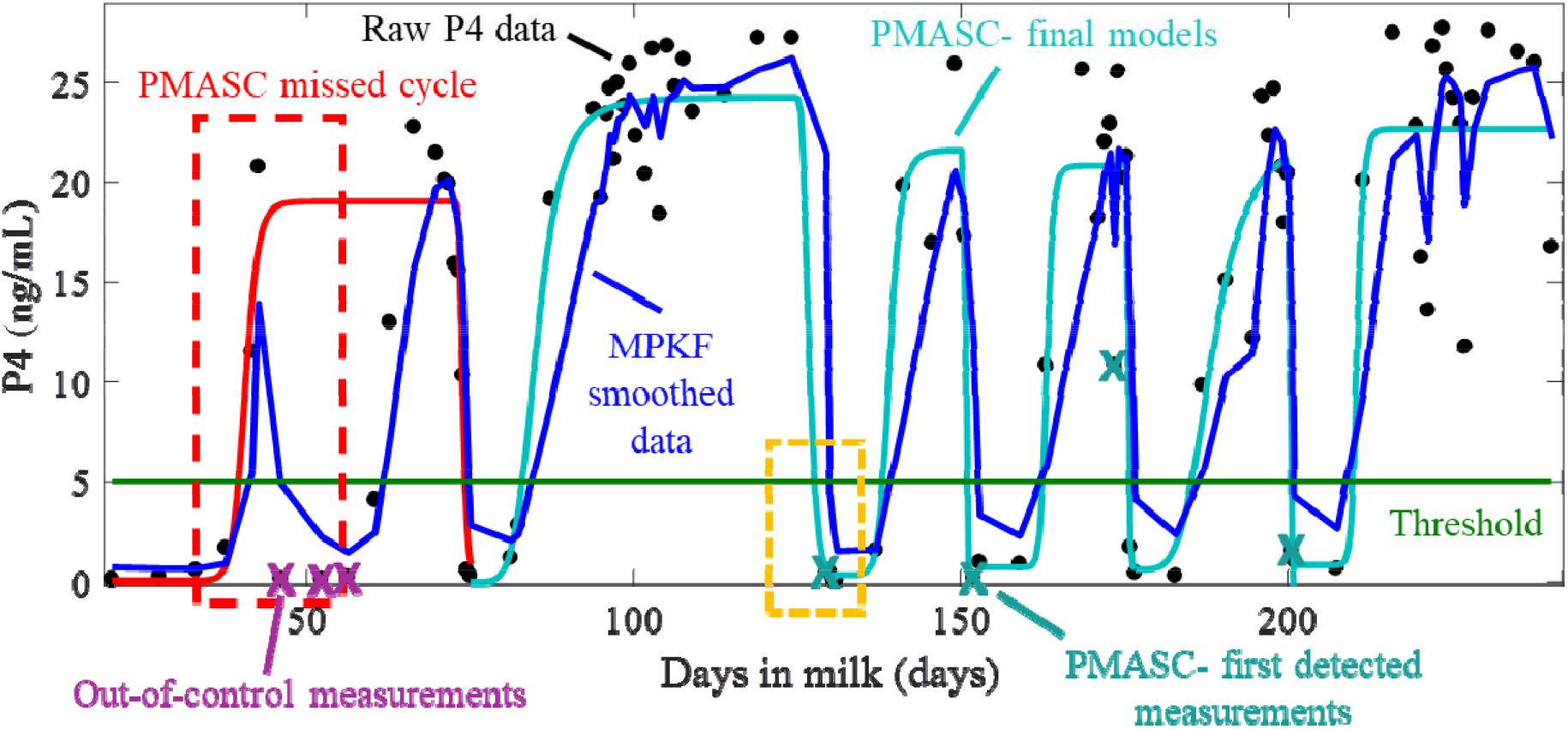
A profile of raw milk progesterone (P4) measurements for one lactation are indicated in black. The light blue crosses are the first out-of-control measurements detected by PMASC, and the light blue lines give final models fitted by PMASC. The first luteolysis was not detected because of an insufficiently high sampling rate (red box), although the measurements pushing the MPKF (indicated in blue) below the threshold (indicated in green) were also out of control for PMASC (indicated with purple cross). The (inconsistent) time-lag between the MPKF dropping below the threshold and actual low measurements of P4 can be seen in the yellow box.

The first out-of-control sample detected by PMASC preceded the MPKF/T alarm on average 20.0±16.1 hours and 2.64 milkings for dataset 1 and 19.8±23.7 hours and 2.2 milkings for dataset 2. The time-delay of the MPKF ensures robustness for outliers and measurement errors, and will correct for unexpected variability. However, contrasting this MPKF smoothed value against a fixed threshold results in alerts for which the timing strongly depends on the absolute P4 values measured and the sampling rate. In cases where the sampling frequency is insufficiently high for timely detection of luteolysis by the MPKF+T method, for example when the sampling or the analysis fails, the alerts of the MPKF+T are less valuable because they arrive too late for timely insemination. However, when these low P4 values are detected by PMASC, information calculated from the model parameters (e.g. time of the inflection point) can be used to interpret these detections and eventually, to adjust the decision process accordingly.

Using the model parameters, the P4 cycle characteristics were calculated. The average cycle length calculated as the time between two successive luteolysis alerts was 27.33±10.0 days, with 56% of the cycles between 19 and 25 days. The average time between the two inflection points of the increasing Hill function and the decreasing Gompertz function was 14.4±10.0 days, which is slightly higher than the time reported in Adriaens et al., (2017), probably because in this previous work only data of normal cycles and profiles were used. The average baseline and maximal P4 concentrations were 0.97±0.77 and 21.5±3.9 ng/mL respectively, and the decrease in P4 at luteolysis was typically fast with a slope of 36.17±24.6 ng/mL per day. The latter means that a drop in P4 from 25 to 2 ng/mL during luteolysis on average takes 14.56 hours, which is in agreement with the values reported by Blavy et al., (2016) and Adriaens et al., (2018b).

In this study, we investigated the performance of PMASC on on-farm measured datasets using less frequent sampling and the LFAI technique, and compared it with the current state of the art being an MPKF+T algorithm. We showed that PMASC, on the condition of a small adjustment in its decision requirements, was able to deal with less samples and the alternative measurement technique, and accordingly, has the potential to work cost-effective in an on-farm setting. More concretely, this study showed that a limited number of samples was sufficient to fit the mathematical model used by PMASC, and did on average not influence the cycle characteristics extracted from it. The non-detected luteolyses were related to cases in which only one sample was taken to confirm estrus, which is rather rare and mainly occurs during the first estrus after calving, in which the maximum P4 concentration remains low. When the sampling rate would be optimized based on information extracted from PMASC (e.g. the time from the previous luteolysis, out-of-control samples etc.) these kinds of artifacts could be avoided. While it would be possible to introduce simple decision rules in the algorithm, the implementation and optimization of the P4 sampling rate based on PMASC was beyond the scope of this study.

## ACKNOWLEDGEMENTS

This work was supported by the Institute for the Promotion of Innovation through Science and Technology in Flanders, Belgium (IWT) [IWT-LA project 110770]. Ines Adriaens is supported by the Fund for Scientific Research (FWO) Flanders, grant number 11ZG916N. We thank Lattec (Hillerod, Denmark) and the commercial dairy farmer who provided us with the second P4 dataset.

